# Deciphering the mechanoresponsive role of β-catenin in Keratoconus epithelium

**DOI:** 10.1101/603738

**Authors:** Chatterjee Amit, Prema Padmanabhan, Janakiraman Narayanan

## Abstract

Keratoconus (KC) a disease with established biomechanical instability of the corneal stroma, is an ideal platform to identify key proteins involved in mechanosensing. This study aims to investigate the possible role of β-catenin as mechanotransducer in KC epithelium. KC patients were graded as mild, moderate or severe using Amsler Krumeich classification. Immunoblotting and tissue immunofluorescence studies were performed on KC epithelium to analyze the expression and localization of β-catenin, E-cadherin, ZO1, α-catenin, Cyclin D1, α-actinin, RhoA, Rac123. Co-immunoprecipitation (Co-IP) of β-catenin followed by mass spectrometry of mild KC epithelium was performed to identify its interacting partners. This was further validated by using epithelial tissues grown on scaffolds of different stiffness. We observed down regulation of E-cadherin, α-catenin, ZO1 and upregulation of Cyclin D1, α-actinin and RhoA in KC corneal epithelium. β-catenin Co-IP from mild KC epithelium identified new interacting partners such as StAR-related lipid transfer protein3, Dynamin-1-like protein, Cardiotrophin-1,Musculin, Basal cell adhesion molecule and Protocadherin Fat 1.β-catenin localization was altered in KC which was validated *in vitro*, using control corneal epithelium grown on different substrate stiffness. β-catenin localization is dependent upon the elastic modulus of the substrate and acts as mechanotransducer by altering its interaction and regulating the barrier function in corneal epithelium.

## Introduction

The mechanical properties of biological tissues are essential for their physiological functions[1] and their regulation plays an important role in driving various physiological and pathological consequences[2]. Such regulation is based on mechanotransductional processes which initiate distinct signalling pathways[3].The cell specific and molecular basis of mechanosensing and the sequence of biochemical reactions that ensue, has been the focus of recent research[4]. Several molecules have been proposed as possible mechanosensors such as integrins, ion channels, G-protein-coupled receptors, and cell–cell junctional proteins[5].

Keratoconus (KC) is a bilateral progressive visually debilitating clinical condition characterized by abnormal thinning and protrusion of the cornea in response to biomechanical instability of the corneal stroma. The etiology of the disease is believed to be multifactorial with genetic and environmental factors being implicated[6]. A decrease in the elastic modulus of the stromal collagen is believed to initiate a cascade of events that results in morphological and biochemical changes that affects the shape and the optical function of the cornea. Morphological changes in the corneal epithelium have been well documented and are considered to be one of the earliest detectable changes observed in KC[7]. Based on the hypothesis that these changes are representative of the mechanotransduction process, we believe that studying the signalling pathway operating in the epithelium of patients diagnosed with KC would offer a unique opportunity to study the molecular basis of mechanotransduction in corneal epithelium.

Wnt/β-catenin pathway which is known to control diverse cell functions[8], respond significantly to extracellular matrix stiffness[9]. The central molecule in Wnt/β-catenin pathway is β-catenin which is an evolutionarily conserved protein and exists in three different subcellular locations: membrane bound, cytosolic, and nuclear[10]. The membrane bound β-catenin has adhesion function whereas its nuclear form plays an important role in transcriptional regulation. In the membrane, β-catenin binds with both E-cadherin and α-catenin which in turn interacts with actin[11].The α-/β-catenin interaction is regulated by small GTPases - RhoA, Rac1 and Cdc42, which in turn regulates actin polymerization[12]. In corneal limbal epithelial stem cells, the nuclear localization of β-catenin is reported to mediate cell proliferation. In contrast, the central corneal epithelium exhibits membrane localization of β-catenin indicating reduced cellular proliferation[13].

The role of β-catenin in KC has not been explored. We hypothesize that β-catenin plays an important role as a mechanotransducer and the detection of its localization could potentially be used for early diagnosis of KC. In this study, we analyzed the localization and expression of β-catenin in KC epithelium. We also studied the expression of its interacting proteins such as E-cadherin, α-catenin, α-actinin, ZO1 (Zonula occludens1). Scanning electron microscopy and Actin staining revealed the structural changes in KC epithelium. β-catenin Co-IP in corneal epithelium of mild KC and normal control identified novel interacting partners. It also explained the role of β-catenin in KC epithelium where canonical Wnt pathway is known to be inactive. Further, the response of β-catenin to different elastic modulus were studied using Polydimethysiloxane (PDMS) and gelatin hydrogel with control corneal epithelium, which further validated our hypothesis that β-catenin does have a role in mechanotransduction.

## Materials and Methods

### Collection of Tissue samples from Patients undergoing Photorefractive corrections and Collagen crosslinking

This study, which conformed to the tenets of the Declaration of Helsinki, was reviewed by the local ethics committee and approved by the institutional review board of Vision Research Foundation, Sankara Nethralaya, Chennai India (Ethics No. 659-2018-P).The tissues used for the study were taken from patients after obtaining their signature and informed consent.

The epithelium of patients undergoing collagen crosslinking for documented KC was used for the study. The epithelium of normal myopic patients undergoing photorefractive keratectomy (PRK) for corrections of their refractive error served as normal control (Ethics No. 489-2015-P). The severity of KC was graded using Amsler-Krumeich classification[14]. The epithelium was collected in DMEM F-12 (Gibco) medium with sodium Pyruvate (Sigma) and transported to the laboratory for further analysis. The tissues were subjected to Immunoblotting, Tissue immunofluorescence, Co-Immunoprecipitation (Co-IP) and primary culture. Concurrently, Polydimethylsiloxane (PDMS) (Sylgard 184 Silicone Elastomer kit) and gelatin (Bovine skin Type B Sigma) hydrogel of different elastic modulus were prepared as described below to serve as substrate for culture of epithelial tissues from normal control.

### Western Blotting

Tissue collected was dissolved in RIPA (Radioimmunoprecipitation assay) buffer(EMD Millipore Cat No-20-188), sonicated in ice for 15s, centrifuged at 13,000 x *g* for 10 min at 4 °C. The supernatant was collected, Proteins were estimated using BCA protein assay kit(Thermo Fisher Cat No-23227). For Western blot, equal concentration of tissue lysates was loaded onto the gel and separated on a sodium dodecyl sulphate polyacrylamide gel (SDS-PAGE) at 100V in electrophoresis buffer (25mM Tris, 190mM Glycine and 0.1% SDS). The proteins were separated and transferred to PVDF membrane (GE Healthcare Cat No-10600023) using semi-dry transblot apparatus (Hoefer) at 1.50 mA/cm^2^. The membrane was then blocked for 1 h at 25 °C in 5% (w/v) non-fat dry milk powder (NFDM) in TBST (20mM Tris-HCl pH 7.5, 150mM NaCl and 0.1% Tween 20). It was washed with TBST, incubated at 4 °C overnight with the following antibodies: E-cadherin, Pan cadherin, α-catenin, β-Catenin, from Cadherin-Catenin antibody sampler kit (Cell signaling and Technology Cat No-9961) β-Catenin for Co-IP(Cell Signaling Technology Cat No-9562S), ZO1, Wnt5ab, RhoA, Rac1/2/3, α-Actinin(Cell signaling and Technology Cat No-13663, 2520T, 21170, 2462, 3134)β-actin, Tenascin C and Cyclin D1 (Santa cruz Cat No-sc47778, sc-20932, sc 246), Syntaxin 3 (Synaptic system 110032). Each antibody was 1000-fold diluted in either 5% (w/v) BSA (Hi Media Cat No-MBO3) or NFDM in TBST. After overnight incubation, the membrane was washed thrice for 5 min with TBST and further incubated in corresponding HRP-conjugated Anti-rabbit and Anti-mouse secondary antibody). The secondary antibodies used were diluted to 10,000-fold in 5% NFDM (w/v) in TBST. After incubation, the membrane was again washed thrice for 5 min with TBST. HRP activity was detected using HRP substrate (Bio-Rad Cat No-1705061) in Bio-Rad Gel Documentation system (Protein Simple).

### Tissue Immunofluorescence

Tissues collected from patients were fixed in 4% Paraformaldehyde and washed with Phosphate buffered saline (PBS).Tissues were permeabilized with 0.5% triton-100 followed by PBS wash. The tissues were then blocked with 1%BSA prior to overnight incubation with primary Antibody. The detection was done using Cy3.5 secondary antibody and counterstained with Hoechest (Thermo fisher Cat No-33342). Further actin fibers of the tissues were stained with 100nM of Phalloidin-488 (Cytoskeleton, inc. Cat No-PHDG1) after permeabilization. The tissues were incubated for 45 minutes in dark and Counter-stained with Hoechest. The tissues were imaged using Zeiss Microscope using 2 different channels.

### Co-Immunoprecipitation followed by Matrix assisted laser desorption/ ionization (MALDI) time-of-flight (TOF)

Control tissues and mild KC tissues were pooled separately after collecting from patients for performing co-immunoprecipitation(Co-IP).Tissue lysates were prepared using RIPA buffer. 400μg lysates, were precleared using 40μl of Protein-A Magnetic beads (New England Biolabs Cat No-S1425S) to remove non-specific components from the lysate. After preclearing 4μg of β-catenin antibody (Cell signaling and Technology, Cat No-9562S) and IgG antibody (Cell signaling and Technology Cat No-2729s) were incubated overnight at 4°C. Further 40μl of Protein A/G Magnetic bead suspension were added and incubated with precleared lysate, agitated for 1h at 4°C. Magnetic field was then applied to pull the beads to the side of the tube and 3 washes were performed using Wash buffer. Further the bead pellet was suspended in 5X sample loading buffer.

### SDS Page and in gel digestion

The immunoprecipitated product was loaded on 10% SDS Polyacrylamide gel. The electrophoresis of the sample was carried out using Tris-Glycine buffer at 100V. After the run was completed the proteins were stained with Coomassie blue stain (Sigma Cat No-G1041). The gel band from IgG Lane, β-catenin IP from control tissue and β-catenin IP from mild KC tissue were excised into 1 mm^3^. The gel was washed with washing solution (50% acetonitrile, 50mM ammonium bicarbonate) until the Coomassie die was completely removed and incubated at room temperature for 15min with gentle agitation. The gel was then dehydrated using 100% acetonitrile(ACN). Then CAN was removed, and the dry gel was kept at room temperature for 10-20min in a vacuum evaporator. The gel was rehydrated for 30min at 56ºC using reduction solution (10mMDTT, 100mM ammonium bicarbonate). Further alkylating solution (50mM iodoacetamide, 100mM ammonium bicarbonate) was added and incubated for 30min in the dark at room temperature. After sequential washing with 25mM NH_4_HCO_3_, 25mM NH_4_HCO_3_/ 50% ACN, and 100% ACN, gel pieces were dried and rehydrated with 12.5ng/mL trypsin (Promega, Madison, WI) solution in 25mM ammonium bicarbonate on ice for 30min. The digestion was continued at 37°C overnight. The tryptic peptides were extracted from gel pieces with extraction solution (60% ACN, 0.1% TFA) and sonicated in ultrasonic water bath for 10min. Further extracted peptides were resuspended in 50% acetonitrile, 0.1% TFA. The samples were spin down and spotted 0.5µL volume on alpha-cyano-4-hydroxycinnamic acid matrix (10 mg/mL in 50% ACN, 0.1% TFA) of MALDI plate. The spots were allowed to dry and further plate was loaded into Voyager.

### Scanning Electron Microscopy

Scanning electron microscopy (FEG-SEM Quanta 400 instrument; FEI), was used to analyze the morphology of epithelial tissue collected after surgery which were fixed with 3.7% glutaraldehyde (Sigma Cat No-G5882) in PBS for 15 min in aluminum-coated coverslip. After washing twice with PBS, the fixed tissues were dehydrated with an ascending sequence of ethanol (40%, 60%, 80%, 96-98%). After evaporation of ethanol, the samples were left to dry at room temperature for 24h on a glass substrate, and then analyzed by SEM after gold–palladium sputtering.

### Preparation and Measurement of Elastic modulus of Polydimethysiloxane gel and Gelatin Hydrogel

Polydimethylsiloxane (PDMS)gel was prepared using polymer and curing agent in a ratio of 1:10 at a temperature of 80ºC for 24h in 24 well plate. Further the PDMS were coated with type-1 collagen (Sigma Cat No-C4243). Hydrogels were prepared using type B gelatin (Sigma G9391). 50mg/ml of gelatin concentration with 0.8 µg/mL of glutaraldehyde as crosslinker were used for preparing hydrogel and incubated at 4°C for 4h for gelation and cross-linking. The cross-linked gelatin hydrogels were immersed in a 50mM glycine aqueous solution under agitation for 1h to block the residual aldehyde groups of glutaraldehyde, followed by two washes with double-distilled water for 1h.

The stiffness of the gels were measured using Microsquischer (Ms.Cell Scale) instrument. The compressive method was employed using 3X3mm plate bound to microbeam. Different forces (500µN, 1000µN, 1500µN) were used for obtaining the force Vs time and force Vs displacement curves. To measure the stiffness, stress (Force/Area) and % strain (tip displacement) were calculated. The graph of stress vs strain was plotted, and linear region was considered to calculate the slope to find the young’s modulus. The rheology (young’s modulus) of hard PDMS gel was measured using MCR301 rheometer (Anton Paar). As a positive control, we used cadaveric stroma along with PDMS. Dynamic oscillary strain sweep experiments were completed on the PDMS/Cadaveric stroma to find the limit of viscoelastic property. The sample size of 1cm radius was used between the compressive plates. The size and thickness of the sample was maintained. The dynamic time sweep was conducted at the frequency of 1 rad s^−1^ and the strain of 0.1% at 25°C. G′ is the storage modulus for measuring the gel-like behavior of a system, whereas G″ is the loss modulus for measuring the sol like behavior of the system.

### Culturing of corneal Epithelial tissue from Patients undergoing Photorefractive corrections

Corneal epithelial cells were cultured in Defined Keratinocyte-Serum Free medium (SFM) (Thermofisher Cat.No-10744019).The tissues collected from control patients was cultivated on type 1 collagen coated 1:10 Polydimethysiloxane (PDMS)of desired elastic modulus and 50mg/ml of Gelatin Hydrogel, in Defined Keratinocyte-SFM medium (Thermofisher Cat.No-10744019). Adherence of the tissues to culture plate was assured by a gravitational force from viscoelastic solution added on top of the tissues (HEALONOVD, Abbott Medical Optics, USA) as reported earlier [15,16]. After 24 hours of incubation tissues were fixed and immunofluorescence were performed.

### Hematoxylin and Eosin staining

Corneal section was deparaffinized in Xylene and then rehydrated in 100% and 95% ethanol. After washing, it was stained with Hematoxylin (Hi-Media Grm 236). The section was differentiated with 1%HCl in 70% ethanol, then washed with water, followed by blueing with weak ammonia. The slide was stained with eosin and then dehydrated slides were mounted.

### Statistical Methods

All experiments were performed at least three independent (n=3) times. However, in case of immunoblotting the number of samples used was mentioned in the legends. The statistical significance of the differences was analyzed by paired student “t” test and one-way analysis of variance (ANOVA) followed by Bonferroni Post-hoc test using Graph Pad software. Asterisks *, **, and *** denote a significance with p-Values <0.05, 0.01, and 0.001 respectively.

## Results

### β-catenin expression and its cellular localization in KC epithelium

Kerataconus is a condition where collagen degradation occurs and decreases the stiffness of stroma. Hence, we investigated the localization of β-catenin throughout the spectrum of disease severity. In control epithelium, β-catenin showed membrane localization whereas in mild moderate and severe KC, β-catenin was observed in the nucleus and cytosol (Fig1A). However, there was no significant difference in the expression of total β-catenin protein (P> 0.05) among KC epithelium compared to control epithelium (Fig1B).Hence, to validate our observation, we studied the expression of β-catenin target gene cyclin D1 that has role in cellular proliferation. In mild KC epithelium. a significant upregulation of cyclin D1(P <0.05) was observed compared to control epithelium (Fig 1C).

**Fig 1.**
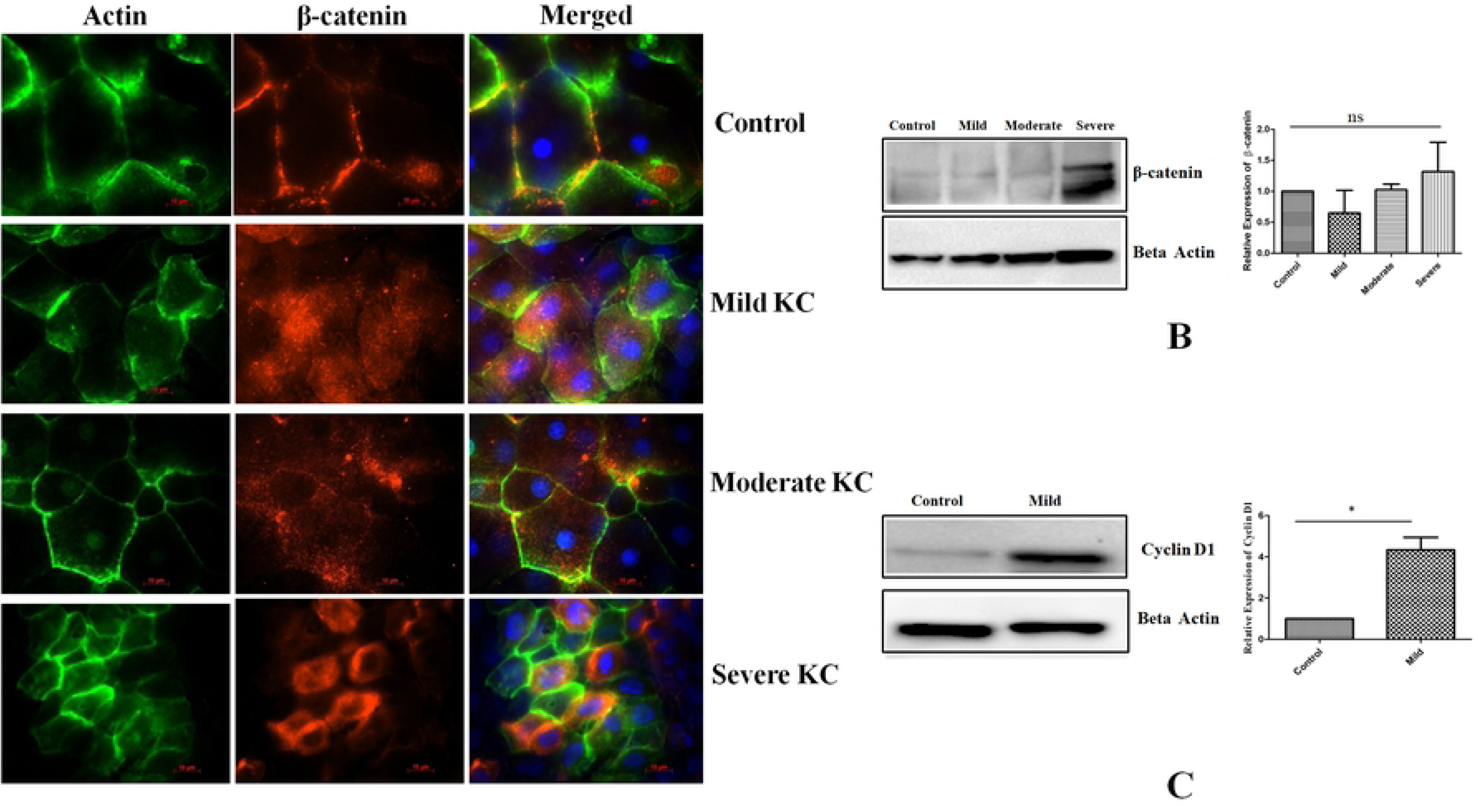
**(A)** Tissue immunofluorescence of β-catenin showing changes in localization in control, mild, moderate and severe keratoconicepithelium (n=3).**(B)**Immunoblotting of β-catenin in control, mild moderate and severe keratoconic epithelium (n=5) One way Anova Asterisks *, **, and *** denote a significance with p-Values <0.05, 0.01, and 0.001 respectively**(C)** Immunoblotting of cyclin D1 in control, and mild keratoconic epithelium (n=5).Student t test *p-Values <0.05.

### Cadherin-Catenin complex in KC epithelium

Since we observed the loss of membrane bound β-cateninin KC epithelium (Fig 1A), we investigated the expression of E cadherin, and α-catenin which are binding partners of β-catenin. A significant down regulation of E cadherin in mild, moderate (P< 0.05), and severe KC epithelium (P< 0.01) was observed compared to control epithelium(Fig 2A).A loss of membrane bound E cadherin was observed in mild KC epithelium tissue (Fig 2B). Additionally, significant upregulation of Pan cadherin levels was observed in KC epithelium compared to control (P< 0.01) (Fig 2C). Since E cadherin-based adhesion is very important for corneal epithelial integrity and is further linked through catenin with actin cytoskeleton, we analyzed the expression of α-actinin which is known crosslinker for actin polymerization. We observed significant upregulation of α-actinin in mild KC epithelium (P< 0.05) suggesting that the KC epithelium is likely to be more migratory in nature (Fig 2D). Furthermore, the expression of α-catenin was significantly downregulated in mild, moderate and severe KC epithelium compared to control (P< 0.001) (Fig 2E). However, we did not observe any significant alteration (P>0.05) in α-catenin and E cadherin expression between different grades (mild, moderate and severe) of KC epithelium. Loss of E cadherin in KC epithelium prompted us to investigate tight junction proteins ZO1 and Claudin1. However, we did not observe significant downregulation of Claudin 1 in mild KC epithelium (P>.05) (Fig 2F). Interestingly, significant loss of ZO1 expression (P< 0.05) and delocalization was observed in the epithelium from mild KC patients (Fig 2G&H).This data suggests that loss of membrane bound β-catenin induces the loss of epithelial integrity in KC epithelium.

**Fig 2.**
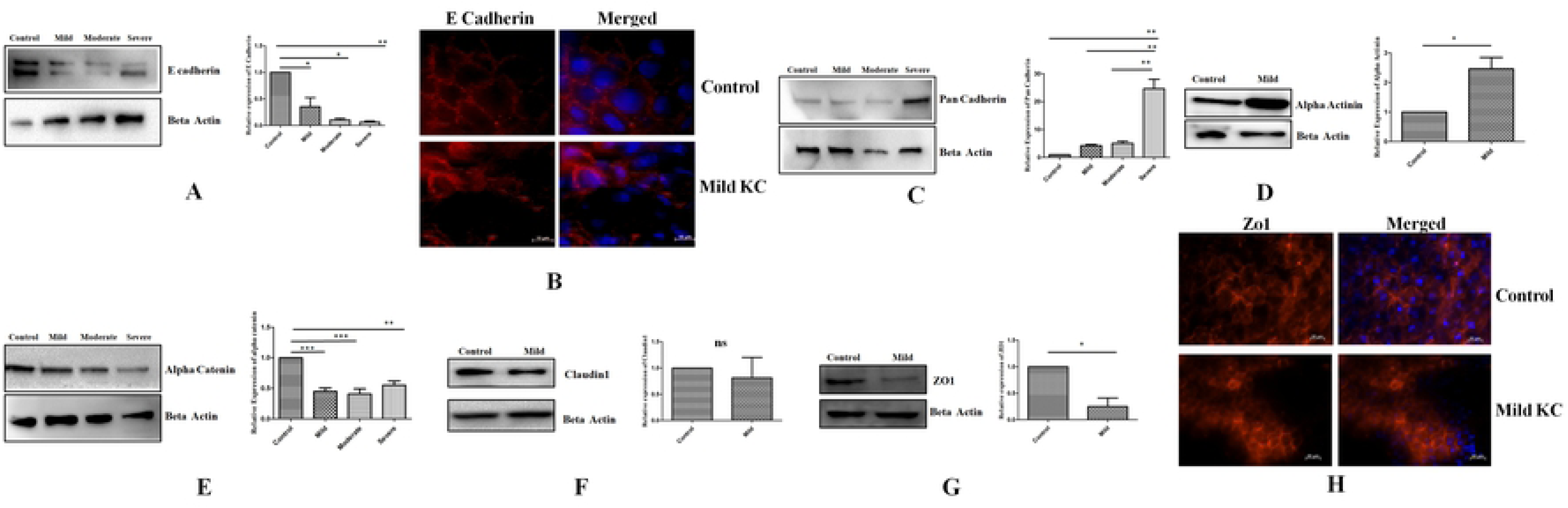
**(A)** Immunoblotting of control, mild, moderate and severe keratoconic epithelium with E cadherin (n=4) **(B)** Tissue Immunofluorescence of E cadherin in control and mild keratoconic epithelium (n=3)**(C)** Immunoblotting of Pan cadherin in control, mild, moderate and severe keratoconic epithelium.(n=3) **(D)** Immunoblotting of α-actinin in control and mild keratoconic epithelium(n=5)Paired two tailed student t test *p-Values <0.05 **(E)** Immunoblotting of α-catenin in control, mild moderate and severe keratoconic epithelium.(n=5) **(F)** Immunoblotting of Claudin1 in control and mild keratoconic epithelium Paired two tailed student t test *p-Values <0.05 (**G&H**) Immunoblotting and Tissue Immunofluorescence of ZO1 in control and mild KC epithelium. Asterisks *, **, and *** denote a significance with p-Values <0.05, 0.01, and 0.001 respectively.

### Morphology and cytoskeleton arrangement of Keratoconic corneal epithelial tissues

The loss of epithelial integrity is known to induce structural changes in the epithelium. Therefore, we assessed the morphological changes using scanning electron microscopy and actin staining. Control epithelium showed uniform thickness (6 to 7 layers) in SEM imaging whereas the severe KC epithelial showed thinning. The basal cells of severe KC epithelium exhibited larger surface than the control epithelium (Fig 3A).The border of KC epithelium was thinner than control epithelium. Cellular morphology and the size of the cells were altered in severe keratoconus compared to the control epithelium(Fig 3A). Grade wise (Mild, Moderate and Severe) analysis of stress fibers in KC epithelium indicated the presence of different types of stress fibers. In mild KC epithelium, stress fibers were located mostly at the membrane of the cells whereas in severe KC epithelium the stress fibers were found across the cells suggesting the changes in the actin polymerization (FigS1). The altered actin polymerization is known to be regulated by Rho family of proteins. Hence, we studied the expression of RhoA, Rac 123 expression in KC epithelium. Expression of RhoA was found to be significantly upregulated (P< 0.05) in severe KC epithelium compared to the control whereas Rac 123 levels were significantly downregulated (P< 0.05)in severe KC epithelium(Fig 3B&C). The altered morphology and expression of actin polymerizing regulating proteins prompted us to analyze the polarity of KC epithelium. STX3 (Syntaxin 3) which localizes to apical plasma membrane and is involved in membrane fusion of apical trafficking pathway, was assessed to understand this phenomenon. The expression of STX3 was not significantly affected in mild KC epithelium whereas in severe KC epithelium it was significantly downregulated (P< 0.05) (Fig 3D).

**Fig 3.**
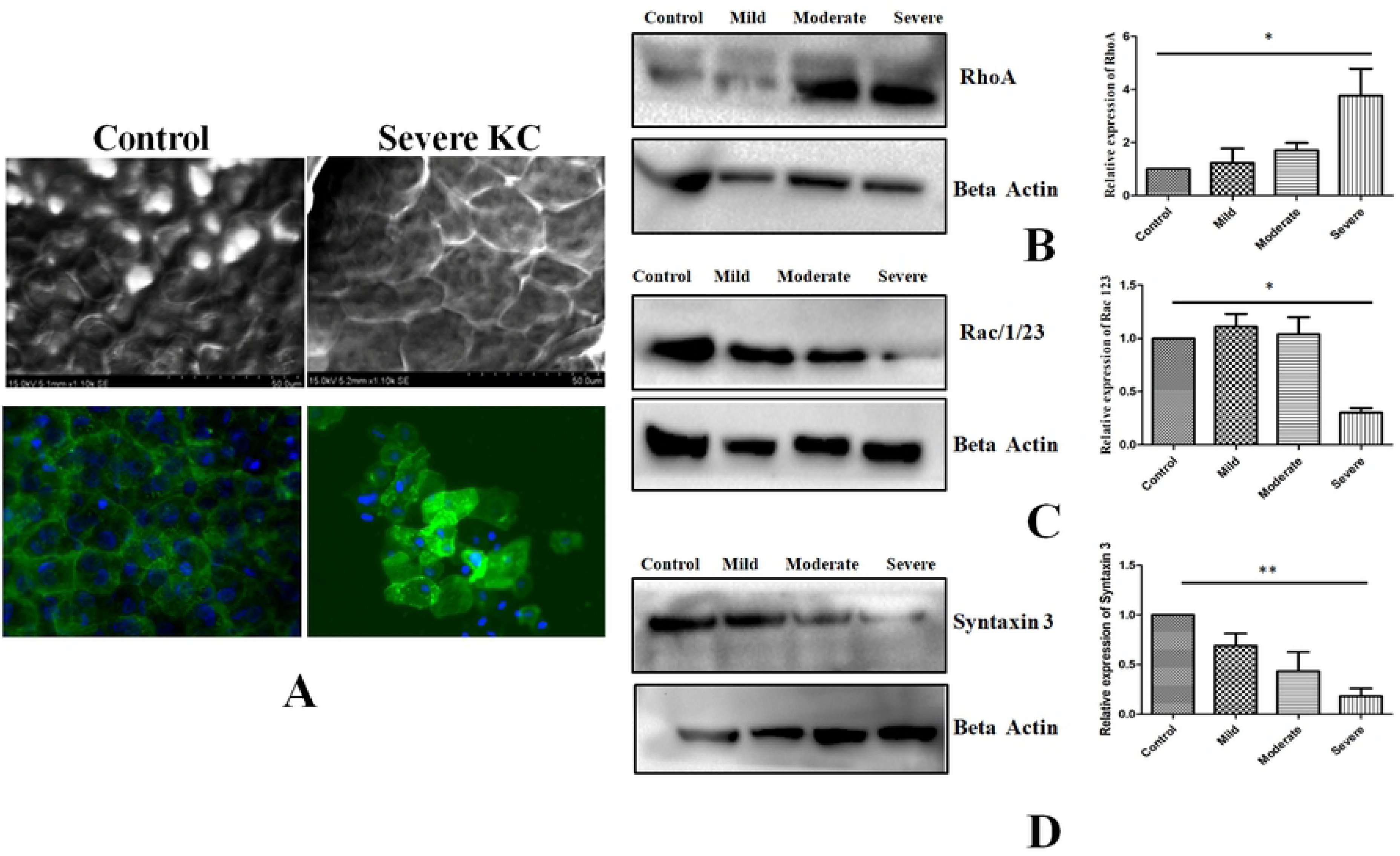
**(A)** Scanning electron microscopy and Phalloidin staining of control epithelium and severe keratoconic epithelium (n=3) **(B)** Immunoblotting of control, mild, moderate and severe keratoconic, RhoA (n=4) **(C)** Immunoblotting of control, mild moderate and severe keratoconic epithelium with Rac123 (n=3) **(D)** Immunoblotting of control mild moderate and severe keratoconic epithelium with STX3 (n=3) Asterisks *, **, and *** denote a significance with p-Values <0.05, 0.01, and 0.001 respectively.

### β-catenin Co-immunoprecipitation and analysis of its interacting partners using mass spectrometry

We next studied the role of Wnt signaling in KC, that mediates either nuclear localization or cytosolic β-catenin degradation by the proteosomes. In KC the expression of β-catenin was unaltered (Fig 1B) indicating lack of proteosomal degradation. Therefore, we next investigated the non-canonical Wnt signaling. We observed that there was no significant difference in the expression level of Wnt5a protein in mild KC epithelium compared to control (Fig4A). The observed stabilization of β-catenin in the cytoplasm may not be due to altered Wnt signaling but could be due to the influence of interacting proteins in this region. Alternatively, the cytoplasmic pool of β-catenin which is destined for ubiquitination and proteasomal degradation can be stabilized by the presence of other interacting proteins in the diseased condition. This prompted us to perform an endogenous Co-IP of β-catenin in control and mild KC epithelium which exhibited undegraded β-catenin, along with IgG control. The presence of β-catenin was observed in the co-immunoprecipitated product (Fig 4B). Then, we looked for the known interacting proteins of β-catenin in this precipitate. This protein interacts with cytoskeletal protein actin which is stabilized by α-catenin and E cadherin proteins. Hence, we investigated whether the loss of membrane bound β-catenin has any effect on these binding partners. Interestingly, we found that there was no loss of interaction between β-catenin and E cadherin whereas interaction between β-catenin and α-catenin was lost in mild KC epithelium (Fig4C).

**Fig 4.**
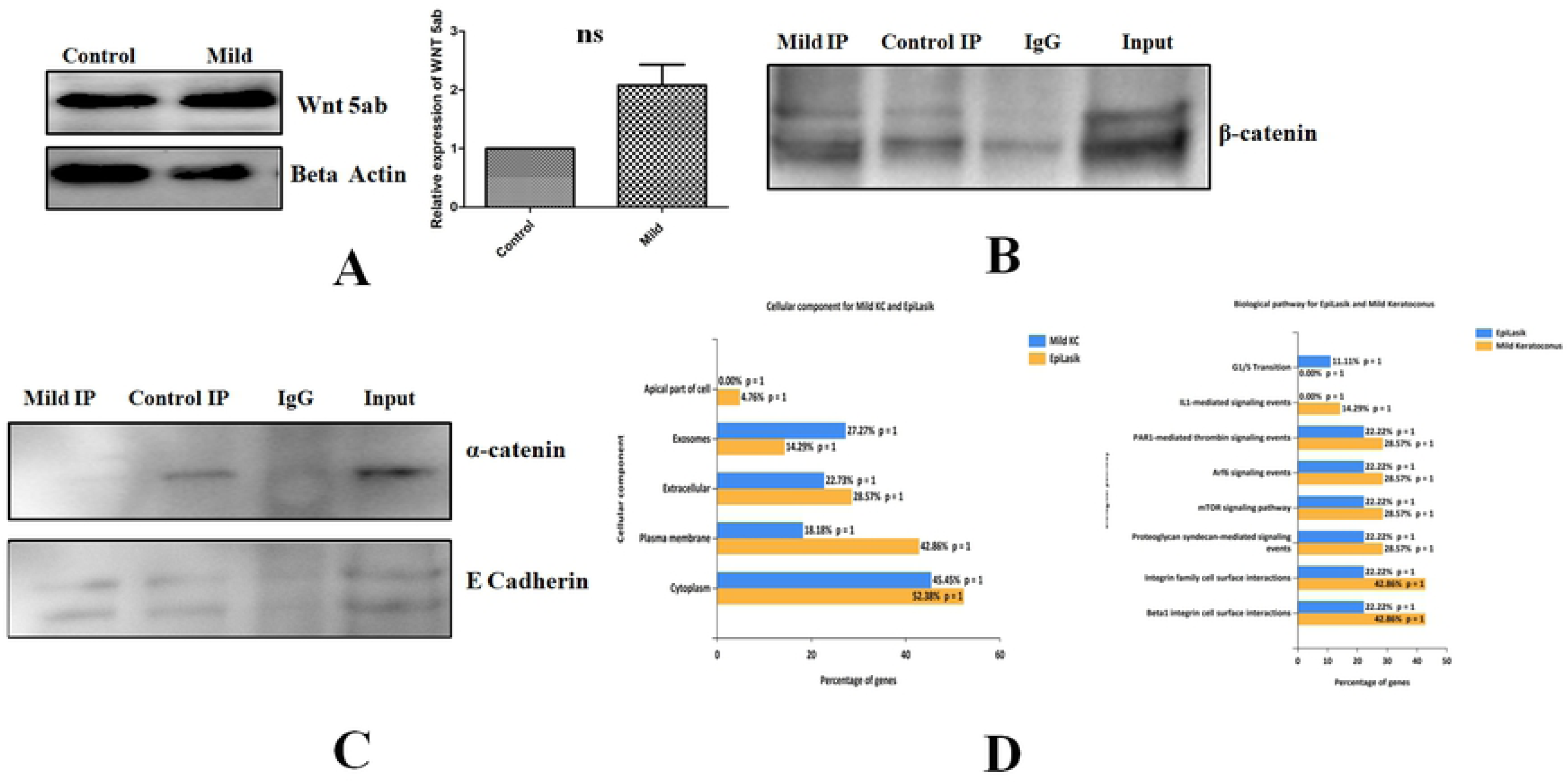
**(A)** Immunoblotting of Wnt5 a in control and mild KC epithelium (n=5). **(B & C)** Immunoblotting of β-catenin, E cadherin and α-catenin for the Co-IP complex in control and mild KC epithelium **(D)** Functional enrichment analysis of the localization and the signaling of proteins which came as a hit in β-catenin Co-IP in control and mild KC epithelium

The Co-IP was performed to capture altered interacting proteins, if any, of β-catenin in the mild stage of the disease as it is stabilized in the cytosol and then compared it to the protein interactions in the control epithelium where this protein has membrane localization. The proteins were separated on SDS gel and were tryptic digested in the gel and mass spectrometry was performed as described in the experimental section to identify novel β-catenin interacting proteins. The proteins were identified from the MS/MS spectra of the peptides (Supplementary S2, MS-MS spectra data) using Mascot server. Endogenous β-catenin from the tissues, the bait protein used for the pull-down assay, was detected in both control and KC tissue lysates. Endogenous Co-IP method contains proteins that are often non-specifically bound to the target protein. Hence, to distinguish these irrelevant background proteins, we devised a negative control using IgG antibody with the same concentration as that of β-catenin antibody. The proteins identified by mass spectrometry in the negative control (IgG immunoprecipitated from control tissues) were subtracted from our downstream data analysis (Table-1) when we compared the protein identifications from the control and the mild KC epithelium. The list of β-catenin interacting proteins in control and mild KC epithelium are listed in (Table 1 and 2). In control epithelium the interacting proteins of β-catenin are Histone Deacetylase 4, Vascular endothelial growth factor receptor1, Sox2 and Rho GTPase-activating protein 21. Some of these proteins are known interacting protein of β-catenin in other cell lines. However, these proteins are novel identifications in corneal epithelium which are lost in mild KC epithelium. The novel interacting proteins observed in the control epithelium are Contactin1, Pappalysin-1, Mimecan which are mostly extracellular matrix regulating and cell surface interacting proteins. The novel β-catenin interacting proteins identified in mild KC epithelium were StAR-related lipid transfer protein 3, Dynamin-1-like protein, Cardiotrophin-1, Musculin, Basal cell adhesion molecule and Protocadherin Fat 1. The known interacting proteins identified are Collagen alpha-1 chain and Laminin subunit alpha-4. Tenascin C was identified in both control and mild KC epithelium. The bioinformatics functional enrichment analysis in the control epithelium identified proteins to delocalized (42.86%) in plasma membrane whereas in mild KC 18.18% of the identified proteins were localized to plasma membrane(Fig4D). Interestingly, Interleukin 1 (IL-1) mediated signaling pathway was enriched by 14.29% in mild KC compared to the control epithelium. Similarly, Integrin signaling pathway enrichment in mild KC was 42.86% compared to the control (Fig4D).

**Table 1:**
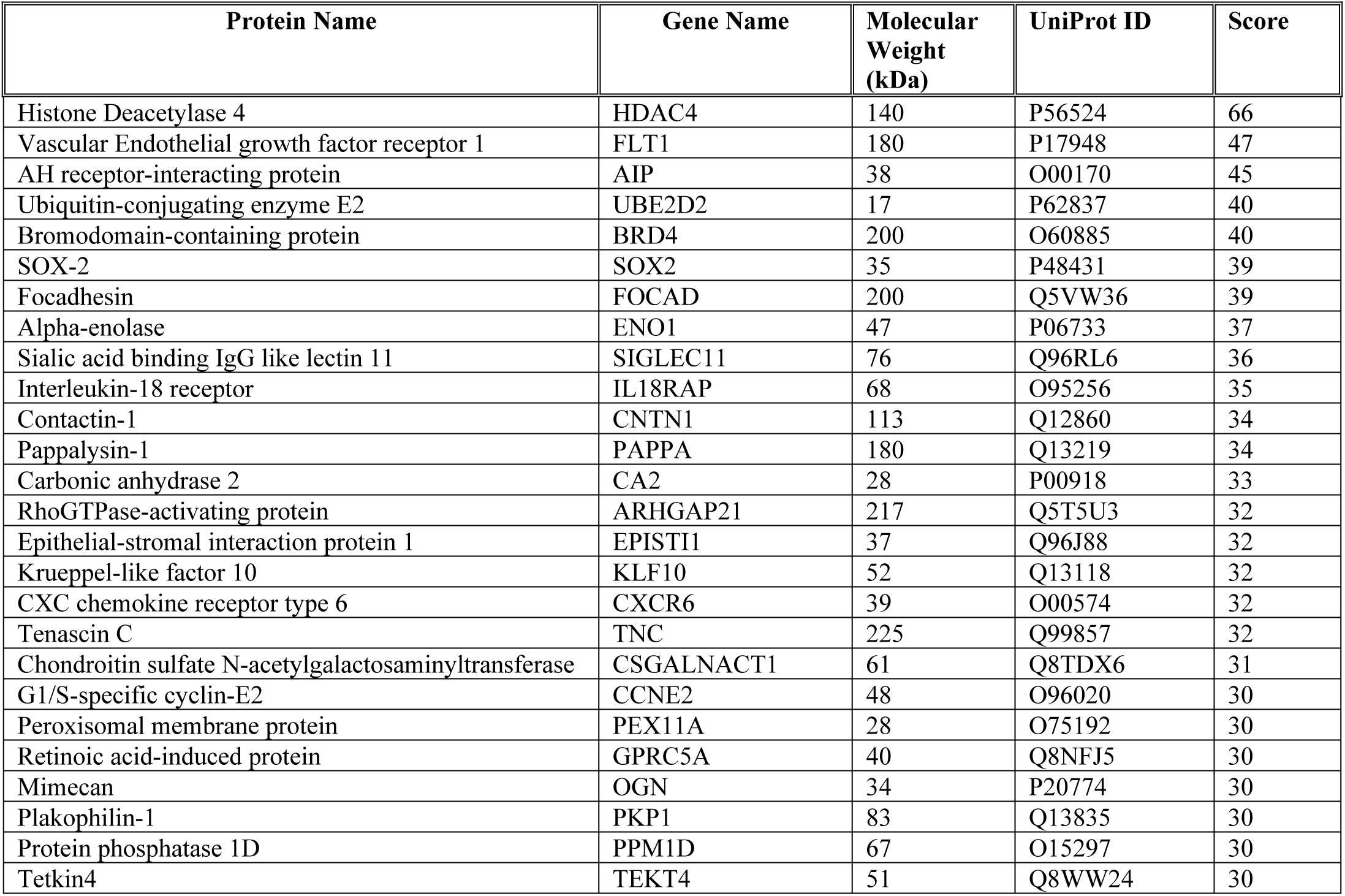
List of β-catenin interacting proteins from control epithelium

**Table 2:** List of β-catenin interacting proteins from mild KC epithelium

| Protein Name | Gene Name | Molecular Wt. (kDa) | UniProt ID | Score |
| --- | --- | --- | --- | --- |
| StAR-related lipid transfer protein 9 | STARD3 | 50 | Q14849 | 51 |
| Protocadherin Fat 3 | FAT1 | 500 | Q14517 | 50 |
| Collagen alpha-1 chain | COL1A1 | 220 | P02452 | 46 |
| Dynamin-1-like protein | DNM1L | 81 | O00429 | 46 |
| Tenascin C | TNC | 225 | Q99857 | 44 |
| Laminin subunit alpha-4 | LAMA4 | 202 | Q16363 | 43 |
| Cilia- and flagella-associated protein 47 | CFAP47 | 361 | Q6ZTR5 | 42 |
| Serine/threonine-protein kinase RIO2 | RIOK2 | 63 | Q9BVS4 | 42 |
| Sperm-associated antigen | SPAG5 | 134 | Q96R06 | 41 |
| DNA-directed RNA polymerase III subunit | POLR3A | 156 | O14802 | 40 |
| A disintegrin and metalloproteinase with thrombospondin motif | ADAMTS5 | 75 | Q9UNA0 | 36 |
| Tumor necrosis factor alpha-induced protein | TNFAIP3 | 82 | P21580 | 36 |
| Anoctamin 6 | ANO6 | 98 | Q4KMQ2 | 35 |
| BBSome-interacting protein | BBIP1 | 10 | A8MTZ0 | 35 |
| Cardiotrophin-1 | CTF1 | 21 | Q16619 | 35 |
| Keratin, type I cuticular | KRT81 | 56 | Q14533 | 35 |
| Probable tubulin polyglutamylase TTLL9 | TTLL9 | 51 | Q3SXZ7 | 34 |
| Activity-regulated cytoskeleton-associated protein | ARC | 27 | Q7LC44 | 33 |
| Musculin | MSC | 22 | O60682 | 32 |
| Aurora kinase A-interacting | AURKAIP1 | 22 | Q9NWT8 | 32 |
| Collagen alpha1 XVI | COL16A1 | 157 | Q07092 | 32 |
| Serine/threonine-protein kinase STK 11 | STK11 | 55 | Q15831 | 32 |
| Basal cell adhesion molecule | BCAM | 67 | P50895 | 31 |
| Oxysterol binding protein 1 | OSBP | 89 | P22059 | 30 |
| RNA polymerase 2 elongation factor | ELL3 | 45 | Q9HB65 | 30 |

### Validation of the Co-IP proteins and β-catenin response to substrate stiffness in control epithelium

Tenascin C that was identified by mass spectrometry in both control and mild KC tissue was validated using Western blotting. We observed this protein expression in both control and KC tissues. Furthermore, we observed upregulation of Tenascin C in mild KC epithelium compared to the control (P<0.05) (Fig5A). Next, we studied a protein that was identified exclusively in KC. We identified collagen alpha chain 1 protein as response to Bowman’s layer breakage and exposure to collagens present exclusively in the stroma. Therefore, we coated the TCP with collagen1 and studied the actin reorganization compared to the non-collagen coated plates. We observed distinct actin staining in corneal epithelial cells grown on collagen matrix compared to the non-coated one(Fig 5B) indicating the effect of modification of ECM on corneal epithelial cells. Similar alterations in ECM could be resulting in KC disease which shows predominant actin staining compared to control(Fig 3B).

**Fig 5.**
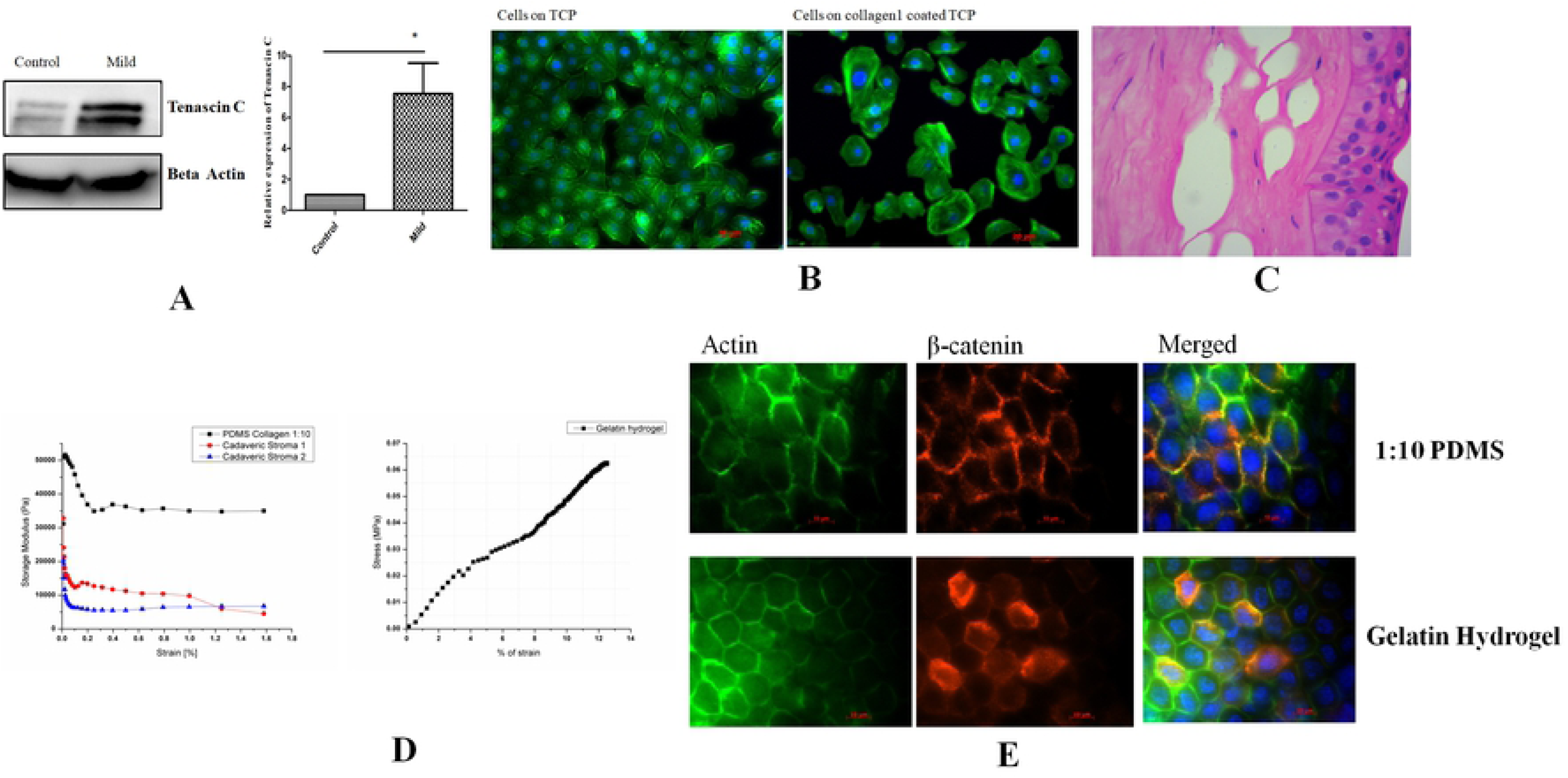
**(A)** Immunoblotting of Tenascin C in control and mild KC epithelium(n=3) **(B)** Phalloidin staining of corneal epithelial cells on plate with and without collagen coating **(C)** Histological section of KC from patients undergoing penetrating Keratoplasty (n=5) **(D)** Elastic modulus measurement of PDMS substrate, cadaveric stroma and Gelatin hydrogel (n=4) **(E)** β-catenin localization of epithelial graft cultured on 1:10 PDMS and Gelatin Hydrogel.

Bowman’s layer is present between epithelium and stroma and is clearly ruptured in KC (Fig5C).We hypothesize that the central epithelium, in the presence of such breaks in the Bowman’s layer, may now sense the corneal stroma which has a different stiffness and activates an altered response. To validate this hypothesis, we prepared PDMS which is 35 kPa and gelatin hydrogel scaffolds whose elastic modulus is 7 kPa (Fig5C). As a control, we measured the elastic modulus of normal cadaveric stroma which was 25±5 kPa which is in agreement to the published reports[17]. β-catenin localization on epithelial graft grown on PDMS showed membrane localization whereas on gelatin hydrogel it showed both membrane and cytoplasmic localization similar to what we had demonstrated in the epithelium of mild KC (Fig5D). Hence, our data suggests that, β-catenin in corneal epithelium responds distinctly to changes in extracellular matrix stiffness.

## Discussion

KC is a bilateral progressive ectatic corneal disease resulting in profound visual impairment. The exact cause and site of origin of the disease remains uncertain. Histopathological changes have been documented in all layers of the cornea except the endothelium. There are several reports in recent times that the corneal epithelium remodels in response to underlying stromal irregularities [18,19] and even suggest that corneal epithelial thickness mapping may be useful in detecting subclinical and early Keratoconus[20,21]. Histological & miRNA studies demonstrate structural & biological changes in corneal epithelium of KC [22]

Substrate stiffness is known to affect cellular behaviour [23]. However, the mechanism underlying mechanosensing in the cornea and the exact signaling pathways involved in the changes of corneal epithelial cells morphology & behavior have not been elucidated yet. β-catenin is a multifunctional protein and its subcellular localization plays a pivotal role in cellular signaling. The membrane bound β-catenin is associated with adhesion function, whereas the nuclear form plays a role in transcriptional regulation [24]. The cytoplasmic pool of β-catenin is highly unstable in the absence of Wnt signal[25]. At the membrane, β-catenin forms complex with E-cadherin and α-catenin which in turn maintains the structural integrity of the epithelial cells. The localization of β-catenin in central corneal epithelium has been reported to be membrane bound [13]. Our results showed that in KC epithelium, the localization of β-catenin has been altered and is cytosolic and nuclear. However, there is no difference in the total β-catenin protein expression. On further investigating the interacting partner of membrane bound β-catenin, we observed significant down regulation of E-cadherin and α-catenin expression in KC corneal epithelium. The role of E-cadherin could be compensated by increased Pan cadherin expression. Additionally, membrane bound E-cadherin was lost indicating loss of cell-cell adhesion. To maintain a polarized epithelium, actin cytoskeleton co-operates with E-cadherin- and integrin-based cell-cell or cell-matrix adhesions[26]. α-actinin which is a major actin filament crosslinker was significantly upregulated in mild KC epithelium. α-actinin upregulation has also been reported in human breast cancer and mammary epithelial cells where it promotes cell migration and induces disorganized acini like structures [27]. The altered KC corneal epithelial structure resulted in the loss of polarity in KC epithelium indicated by lowered syntaxin 3 expression. Furthermore, loss of the epithelial polarity could lead to altered tight junction structure and function, ZO1 tight junction proteins was significantly downregulated in KC epithelium. The loss of both E-cadherin and ZO1 could be associated with the observed change in morphology of KC epithelium which has altered cellular adhesions [7]. The altered cell-cell adhesion might have caused differential stress fibers localization in the KC epithelium. In the mild KC epithelium, stress fibers were located mostly at the membrane of the cells whereas in severe KC epithelium the stress fibers were found across the cells. This could be due to changes in the actin polymerization. The alteration in stress fiber formation and actin polymerization are regulated by RhoA, and Rac,[28]. Hence, on analyzing the expression of these protein we found that the expression pattern matches with the phenotype of the epithelium. The observation from our studies suggested that loss of membrane bound β-catenin may be reason for the loss of adherence junction proteins which in turn altered the cellular morphology of KC epithelium. Consistent with our studies, an altered cellular morphology of reduced basal epithelial cell density in moderate and severe keratoconus and greater average basal epithelial cell diameter were observed in keratoconic patients [7,29].

The localization of β-catenin is also dependent on Wnt signalling [30]. However, this protein may not be involved in disease progression of KC as this protein was unaltered in the diseased condition. β-catenin is a central molecule in Wnt signaling pathway and the transcriptional activity of β-catenin is also modulated by direct regulation of its subcellular localization by a variety of interacting partners [31]. In mild KC epithelium in the absence of the Wnt signaling, the loss of interaction between β-catenin and α-catenin might lead to lowered ubiquitylation and stabilization of this protein. Similarly, the ubiquitylation of native β-catenin was strongly inhibited in α-catenin depleted cells leading to stabilization of the protein[32]. Additionally, the known interacting proteins Collagen alpha-1 chain and Laminin subunit alpha-4 could also alter the localization of β-catenin. Furthermore, in response to collagen 1 coating, actin reorganization was altered in corneal epithelial cells. β-catenin localization was altered in response to matrix stiffness (PDMS to gelatin hydrogel) mimicking the scenario in KC. Additionally, although Tenascin C was found in both control and mild KC epithelium, it was upregulated in KC due to the presence of the β-catenin in the cytosol and nucleus. Tenascin C has an antiadhesive activity and has been reported to be regulated by β-catenin[33]. Therefore, we summaries that β-catenin might sense the altered stiffness in KC when the Bowman’s layer is ruptured. It is established that various layers of cornea exhibit different elastic moduli [17] and as the disease progresses the epithelium is exposed to matrix of different elastic moduli. Therefore, we can conclude that β-catenin could be a putative mechanotransducer and its localization could potentially be used for early diagnosis of KC. We also observed novel interacting partners for β-catenin in KC epithelium such as StAR-related lipid transfer protein 3, Dynamin-1-like protein, Cardiotrophin-1, Musculin, Basal cell adhesion molecule and Protocadherin Fat 1. The role of these binding proteins when β-catenin alters localization in KC needs further investigations.

## Acknowledgement

JN and AC acknowledge the Department of Biotechnology (DBT), Government of India, for funding (BT/PR14690/MED/32/496/2015) and fellowship.We are grateful to Dr. Jyotirmay Biswas for the histopathology. We also thank Prof. James Chodosh (Massachusetts Eye and Ear Boston, Massachusetts, United States) for the kind gift of corneal epithelial cells. We thank Dr. Sailaja V. Elchuri, Department of Nanobiotechnology, Vision Research Foundation, Sankara Nethralaya campus, for critically reviewing the manuscript

## Authors contribution

AC, JN and PP conceived the idea. AC, JN and PP designed the experiments. JN received the funding. AC carried out the experiments, PP provided clinical samples. AC, JN and PP analysed the data. AC written the manuscript. JN, AC, PP reviewed the manuscript.

## Supplementary Information

**Supplementary S1:** Actin staining of control and KC tissues

**Supplementary S2:** MS-MS spectra of β-catenin Co IP from control and mild KC

